# Star allele search: a pharmacogenetic annotation database and user-friendly search tool of publicly available 1000 Genomes Project biospecimens

**DOI:** 10.1101/2023.07.27.550362

**Authors:** N. Gharani, G. Calendo, D. Kusic, J. Madzo, L. Scheinfeldt

**Affiliations:** Coriell Institute for Medical Research, 403 Haddon Ave, Camden, NJ 08103, USA; Gharani Consulting Limited, 272 Regents Park Road, London, N3 3HN, UK

**Keywords:** Pharmacogenetic, pharmacogenomic, reference material

## Abstract

Here we describe a new public pharmacogenetic (PGx) annotation database of a large (n=3202) and diverse biospecimen collection of 1000 Genomes Project cell lines and DNAs. The database is searchable with a user friendly, web-based tool (www.coriell.org/StarAllele/Search). This resource leverages existing whole genome sequencing data and PharmVar annotations to characterize *alleles for each biospecimen in the collection. This new tool is designed to facilitate in vitro functional characterization of *allele haplotypes and diplotypes as well as support clinical PGx assay development, validation, and implementation.

## Background

Pharmacogenomics (PGx) holds the potential to improve medication management by increasing efficacy and by reducing toxicity [1-7]. Translating pharmacogenomic research into clinical care, however, requires a robust inter-disciplinary infrastructure [8, 9]. Characterizing the full range of functionally relevant human pharmacogenetic variation is limited by the documented underrepresentation of many communities living in the United States and around the world [10-16], and this effort would benefit from a large and diverse collection of publicly available and well-characterized cell lines. Such a resource would facilitate a more comprehensive understanding of pharmacogene variation and in vitro drug response [17-23]. Moreover, a well-characterized and diverse set of publicly available and renewable DNA samples would benefit the clinical communities that require positive and negative controls for assay development, validation, implementation and proficiency testing for robust PGx testing.

The Genetic Reference and Testing Materials Coordination Program (GeT-RM) has used a variety of clinical testing methods to characterize lymphoblastoid cell line (LCL) DNAs for 28 pharmacogenes [24], and more recently has incorporated next generation sequencing data for the characterization of *CYP2D6* [25] (n=179), as well as *CYP2C8, CYP2C9* and *CYP2C19* (n=137) [26]. Here we describe a complementary PGx annotation resource that includes a significantly larger set (n=3,202) of renewable and publicly available 1000 Genomes Project LCLs and DNAs available through the NHGRI Sample Repository for Human Genetic Research (https://catalog.coriell.org/1/NHGRI) and the NIGMS Human Genetic Cell Repository (https://catalog.coriell.org/1/NIGMS). This new annotation resource leverages 30x whole genome sequencing (WGS) data [27], is downloadable (**Table S1**) and may be searched with a user-friendly, web-based tool (www.coriell.org/StarAllele/Search).

### Construction and content

**Table 1** includes a summary of all of the publicly available 1000 Genomes Project biospecimens included in the star allele annotation database. The majority of the samples (n=3023) are available through the NHGRI Sample Repository for Human Genetic Research (https://catalog.coriell.org/1/NHGRI), and the collection of Utah Residents (CEPH) with Northern and Western European Ancestry biospecimens (n=179) are available through the NIGMS Human Genetic Cell Repository (https://catalog.coriell.org/1/NIGMS). **Table S1** includes each individual NHGRI Sample Repository for Human Genetic Research and NIGMS Human Genetic Cell Repository identifier for all of the 1000 Genomes Project biospecimens annotated in the star allele annotation database.

**Table 1.** List of Included Populations.

| **Population Descriptor** | **N samples with NYGC WGS data** |
| --- | --- |
| African Caribbean in Barbados | 116 |
| Bengali in Bangladesh | 131 |
| British from England and Scotland | 91 |
| Chinese Dai in Xishuangbanna, China | 93 |
| Colombian in Medellin, Colombia | 132 |
| Esan in Nigeria | 149 |
| Finnish in Finland | 99 |
| Gambian in Western Division - Mandinka | 178 |
| Han Chinese South, China | 163 |
| Iberian Populations in Spain | 157 |
| Indian Telugu in the UK | 107 |
| Kinh in Ho Chi Minh City, Vietnam | 122 |
| Mende in Sierra Leone | 99 |
| Peruvian in Lima, Peru | 122 |
| Puerto Rican in Puerto Rico | 139 |
| Punjabi in Lahore, Pakistan | 146 |
| Sri Lankan Tamil in the UK | 114 |
| African Ancestry in Southwest USA | 74 |
| Gujarati Indians in Houston, Texas, USA | 103 |
| Han Chinese in Beijing, China | 103 |
| Japanese in Tokyo, Japan | 104 |
| Luhya in Webuye, Kenya | 99 |
| Mexican Ancestry in Los Angeles, California, USA | 97 |
| Toscani in Italia | 107 |
| Yoruba in Ibadan, Nigeria | 178 |
| Utah Residents (CEPH) with Northern and Western European Ancestry* | 179 |
| **Total Samples** | **3202** |

\* samples available through the NIGMS Human Genetic Cell Repository

We leveraged existing publicly available 30x coverage WGS data generated from 3202 samples and phased by the New York Genome Center (NYGC) [27]. The detailed description of the data collection and analysis can be found in Byrska-Bishop et al. [27]. Briefly, 3,202 samples from the 1000 Genomes Project collection were selected for inclusion [27] in the WGS data collection (**Table 1**). The sample set includes 2,504 unrelated individuals as well as 698 relatives (that together complete 602 complete trios) [27], and the WGS data were collected with an Illumina NovaSeq 6000 System [27]. The raw WGS data were aligned to the GRCh38 reference genome, and variant calling was performed with GATK [27, 28]. The WGS variant information was additional phased into haplotypes; autosomal single nucleotide variants (SNVs) and insertion / deletions (INDELs) were statistically phased using SHAPEIT-duoHMM with pedigree-based correction [27, 29, 30].

We used the phased NYGC WGS VCF files [27] identified through the www.internationalgenome.org website (https://www.internationalgenome.org/data-portal/data-collection/30x-grch38), and accessed from the following publicly available FTP site (https://ftp.1000genomes.ebi.ac.uk/vol1/ftp/data_collections/1000G_2504_high_coverage/working/20220422_3202_phased_SNV_INDEL_SV/), which were last modified on 2022-11-14 08:33. This dataset was filtered prior to phasing according to the following criteria as described in the NYGC README file: 1) missing genotype rate >5%; 2) Hardy Weinberg test P-Value >1e-10; Mendelian error rate ≤5%; and 3) minor allele count ≥2. We did not perform any additional data post processing. We compared the variants in the phased VCF files against PGx annotations for 12 of the 13 pharmacogenes annotated in PharmVar [31-33] version 5.2.13 using ursaPGx [34]which implements Cyrius for *CYP2D6* calling using raw WGS BAM files [35]. The detailed description of the ursaPGx annotation can be found here [34]. Briefly, for each pharmacogene, the star allele defining variants according to PharmVar are extracted from the phased VCF file, and the annotation is assigned when all star allele defining variants are present for a given VCF haplotype. In cases where no complete match between the phased haplotype and the PharmVar star allele occurs, the haplotype is annotated as ambiguous.

### Utility and discussion

Here we describe a new public PGx annotation database with a user friendly, web-based search tool of associated lymphoblastoid cell line and DNA biospecimens. This new resource complements existing databases generated by GeT-RM; while GeT-RM works directly with clinical laboratories to develop robust PGx annotated biospecimens designed to serve as reference materials for genetic testing, this effort is extremely involved and not easily scalable to larger collections of biospecimens. This new annotation database therefore offers a slightly less robust characterization of a significantly larger collection of diverse biospecimens (**Table 1**) to support PGx related research efforts and to serve as a starting point for clinical testing communities to identify potentially relevant reference materials for their testing needs.

To assess the quality and accuracy of the PGx annotation database, we compared overlapping samples that were already characterized by GeT-RM using next-generation sequencing data, which is available for *CYP2C8, CYP2C9, CYP2C19*, and *CYP2D6* [25, 26]. In total, we identified 87 overlapping samples between GeT-RM and the current annotation dataset [25, 26]. We found 100%, 99% and 97% concordance, respectively between our annotation and the GeT-RM NGS consensus annotation for *CYP2C8, CYP2C19, and CYP2C9* [26], and we found 94% concordance between our annotation and the GeT-RM NGS consensus annotation for *CYP2D6* [25].

Our *CYP2C19* comparison identified a discrepancy for a single sample (NA19122). The GeT-RM NGS consensus is *2/*35 [26], while our annotation is *2|*Amb. We note that as described above, our approach requires a complete match between a given phased haplotype and all of the PharmVar defining variants for a given star allele. For NA19122, the first haplotype included all of the variants required to annotate *2 (non-reference alleles for rs12769205, rs4244285, rs3758581), consistent with GeT-RM [26]; however, the second haplotype in our phased VCF file includes both variants required to annotate *35 (non-reference alleles for rs12769205, rs3758581) as well as a non-reference allele at rs17882687, which in our approach precludes it from an unambiguous call of *35 or *15.

We identified discordant *CYP2C9* star allele annotations for three samples. Our approach annotated two samples (NA19143, NA19213) as *1|*1 while the GeT-RM NGS consensus is *1/*6 [26]. This discrepancy is due to the limitation of the WGS phased VCF file we used which unfortunately does not contain rs9332131, which is the single base deletion that defines *6. We annotated the third discordant sample (HG01190) to be *61|*1, whereas the GeT-RM NGS consensus is *2/*61 [26]. We believe this difference is due to differences in variant calling and phasing approaches. In the phased VCF we used, this sample is heterozygous for both variants required to annotate *61 (rs1799853 and rs202201137), and both of these variants occur on the first haplotype of the sample. Here we also note that while the consensus annotation is *2/*61, a minority of the groups participating in the study annotated this sample as *1/*61 [26].

For *CYP2D6*, we identified a perfect match for 82 overlapping samples, with an additional two samples (NA07000, NA19143) concordant between the ursaPGx implementation of Cyrius and the tentative GeT-RM assignment designated with parentheses [25]. For NA18519, the ursaPGx implementation of Cyrius annotated *106/*29, while GeT-RM annotated *1/*29. As far as we can tell from the detail included in Table S2 of the publication’s supplementary materials [25], the *106 defining variant (rs28371733) was not included in the NA18519 annotation assessment; *106 was not detected by the assays used for the full set of included samples (n=179), including genotyping, PharmacoScan, iPLEX V1.1, CYP2D6 V1.1, a custom panel, and VeriDose, but rather sequencing (Sanger, NGS or SMRT) appears to have been used for a subset of 50 samples that did not include NA18519. The ursaPGx implementation of Cyrius was not able to resolve a single diplotype for NA19908, but rather annotated two possible diplotypes (*1/*46;*43/*45) due to short read phase uncertainty [35]. GeT-RM utilized long-range PCR to resolve the phase uncertainty and correct diplotype for this sample [25]. Similarly, the ursaPGx implementation of Cyrius was not able to fully resolve the diplotype for NA18565 using short read WGS data beyond *36/*36+*10, while GeT-RM was able to fully resolve the diplotype to *10/*36x2 (one *10 allele and a second allele with two copies of *36).

In addition to the GeT-RM annotation benchmarking, we compared PGx annotation using the newest NYGC 30x WGS dataset available [27] against the older Phase 3 10x coverage WGS dataset available for a subset of 2504 unrelated individuals [36] for *CYP2C9* and *CYP2C19*. Several *allele-defining variants were present in the Phase 3 10x dataset but absent from the NYGC phased 30x VCF files (**Table S2**). In particular *CYP2C9* *6 (rs9332131, A deletion), *7 (rs67807361, A allele), *16 (rs72558192, G allele), *33 (rs200183364, A allele), *36 (rs114071557, G allele), *45 (rs199523631, T allele), *63 (rs141489852, A allele), *68 (rs542577750 A allele), and *73 (rs17847037, T allele) and *CYP2C19* *16 (rs192154563, T allele), *24 (rs118203757, A allele), and *30 (rs145328984, T allele).

In total, star allele search includes 956 diplotypes across 14 pharmacogenes (**Table 2, Table S1**), excluding diplotypes with one or two ambiguous (i.e., Amb) allele calls. Each unique diplotype and associated diplotype frequency in the database is detailed in **Table S3**, and each unique *allele haplotype and associated haplotype allele frequency in the database is detailed in **Table S4**. To determine the contribution of the larger sample set included in the database, we identified 3, 3, 7, and 10 new *alleles, respectively in the dataset relative to GeT-RM [25, 26] for *CYP2C19* (*11, *22, *34), *CYP2C8* (*6, *11, *14), *CYP2C9* (*12, *13, *14, *29, *31, *44, *66), and *CYP2D6* (*27, *32, *34, *49, *84, *86, *117, *121, *125, *139) (**Figure 1, Table S5**). We performed a similar comparison for unique pairs of *alleles (diplotype combinations). We chose to conservatively exclude ambiguous calls, copy number variants and complex *CYP2D6* structural variants and identified 12, 17, 23, and 132 new diplotypes, respectively for *CYP2C8, CYP2C19, CYP2C9*, and *CYP2D6* (**Figure 1, Table S5**).

**Table 2.**
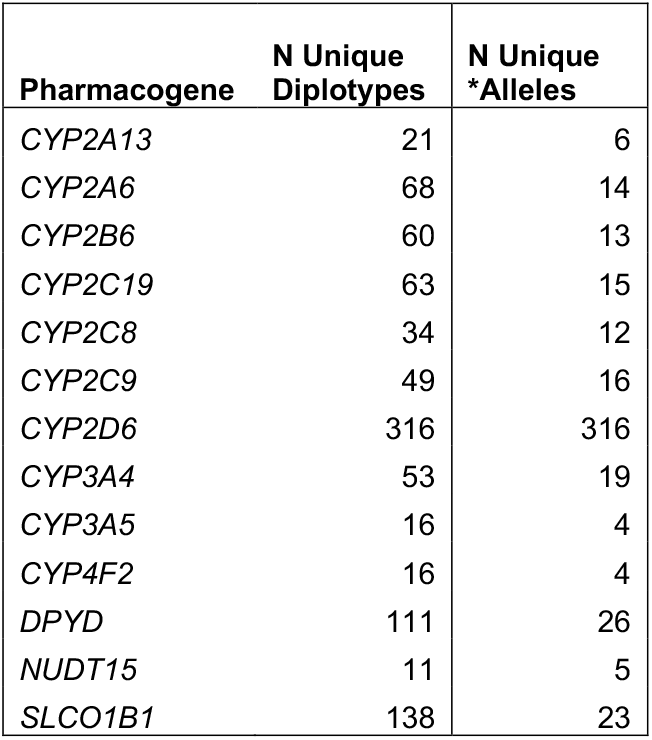
List of PharmVar annotated pharmacogenes, number of diplotypes and *alleles included in database.

| **Pharmacogene** | **N Unique**<br>**Diplotypes** | **N Unique**<br>**\*Alleles** |
| --- | --- | --- |
| *CYP2A13* | 21 | 6 |
| *CYP2A6* | 68 | 14 |
| *CYP2B6* | 60 | 13 |
| *CYP2C19* | 63 | 15 |
| *CYP2C8* | 34 | 12 |
| *CYP2C9* | 49 | 16 |
| *CYP2D6* | 316 | 316 |
| *CYP3A4* | 53 | 19 |
| *CYP3A5* | 16 | 4 |
| *CYP4F2* | 16 | 4 |
| *DPYD* | 111 | 26 |
| *NUDT15* | 11 | 5 |
| *SLCO1B1* | 138 | 23 |

**Figure 1.**
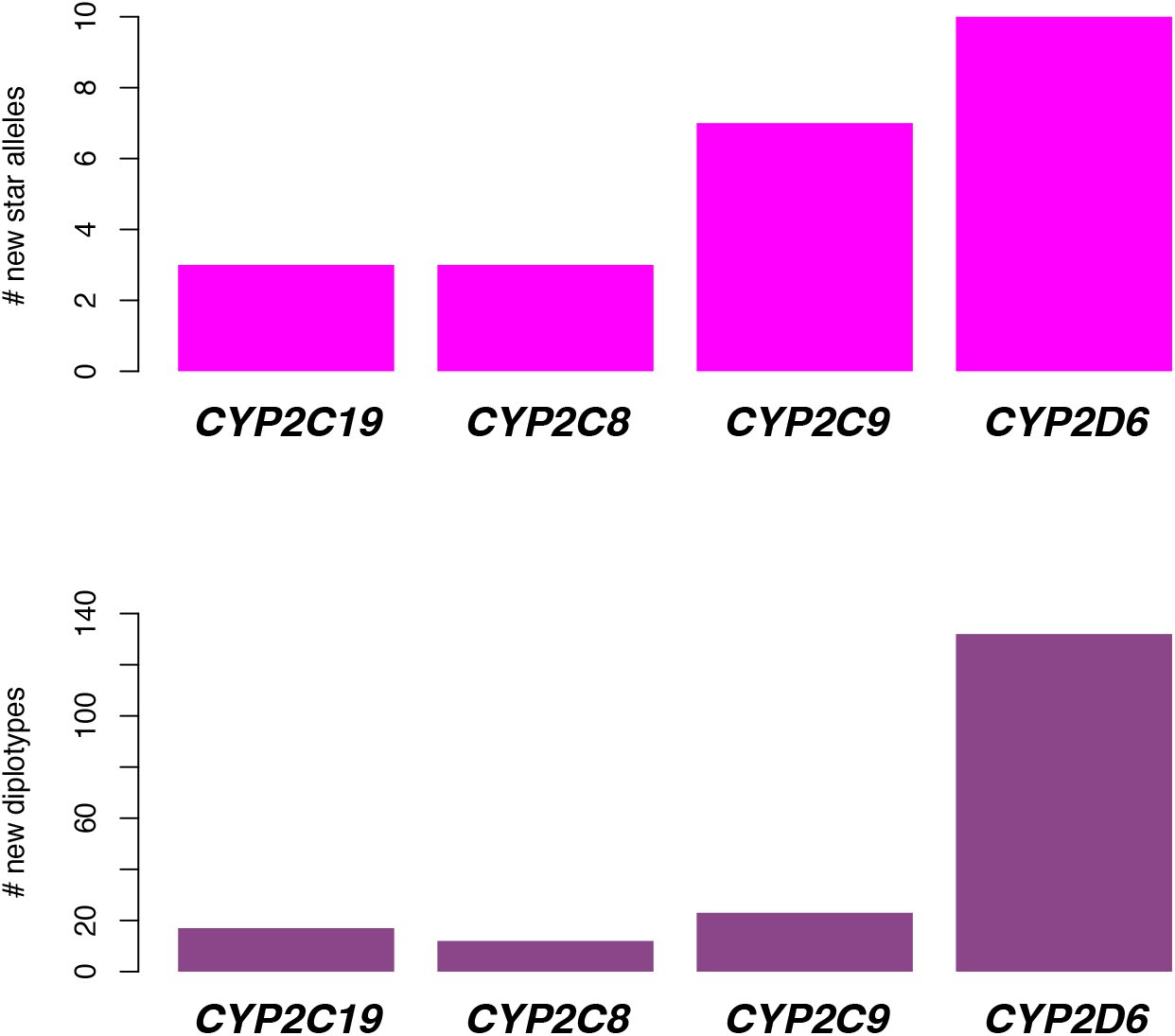
New *alleles and diplotypes identified in sample set (n=3,202) relative to GeT-RM. The top panel of Figure 1 displays the total number of new star alleles (Y-axis) for *CYP2C19* (n=3), *CYP2C8* (n=3), *CYP2C9* (n=7) *and CYP2D6* (n=10), respectively along the X-axis in magenta. The bottom panel of Figure 1 displays the total number of new diplotypes (Y-axis) for *CYP2C19* (n=17), *CYP2C8* (n=12), *CYP2C9* (n=23) *and CYP2D6* (n=132), respectively along the X-axis in purple.

This new star allele annotated biospecimen database is of use for a wide range of applications. For example, researchers interested in functionally characterizing *alleles of interest can use the resource to choose LCLs with the most relevant diplotype combinations; researchers interested in developing new PGx assays can use the resource to benchmark performance; and clinical laboratories can use the resource to minimize the number of positive and negative control DNAs needed for a given PGx test.

We have additionally developed Star Allele Search (**Figure 2**), which is a web-based search tool of the new PGx biospecimen annotation database to facilitate these types of research and clinical applications. In addition to this new database and search tool, users can choose to search existing public data one variant at a time, up to one hundred variants at a time, or by gene (https://www.coriell.org/SNPSearch/WGS; [27]). Users can also search gene expression data collected from a subset of the *allele annotated LCLs (n=462) (http://omicdata.coriell.org/geuv-expression-browser/; [37]).

**Figure 2.**
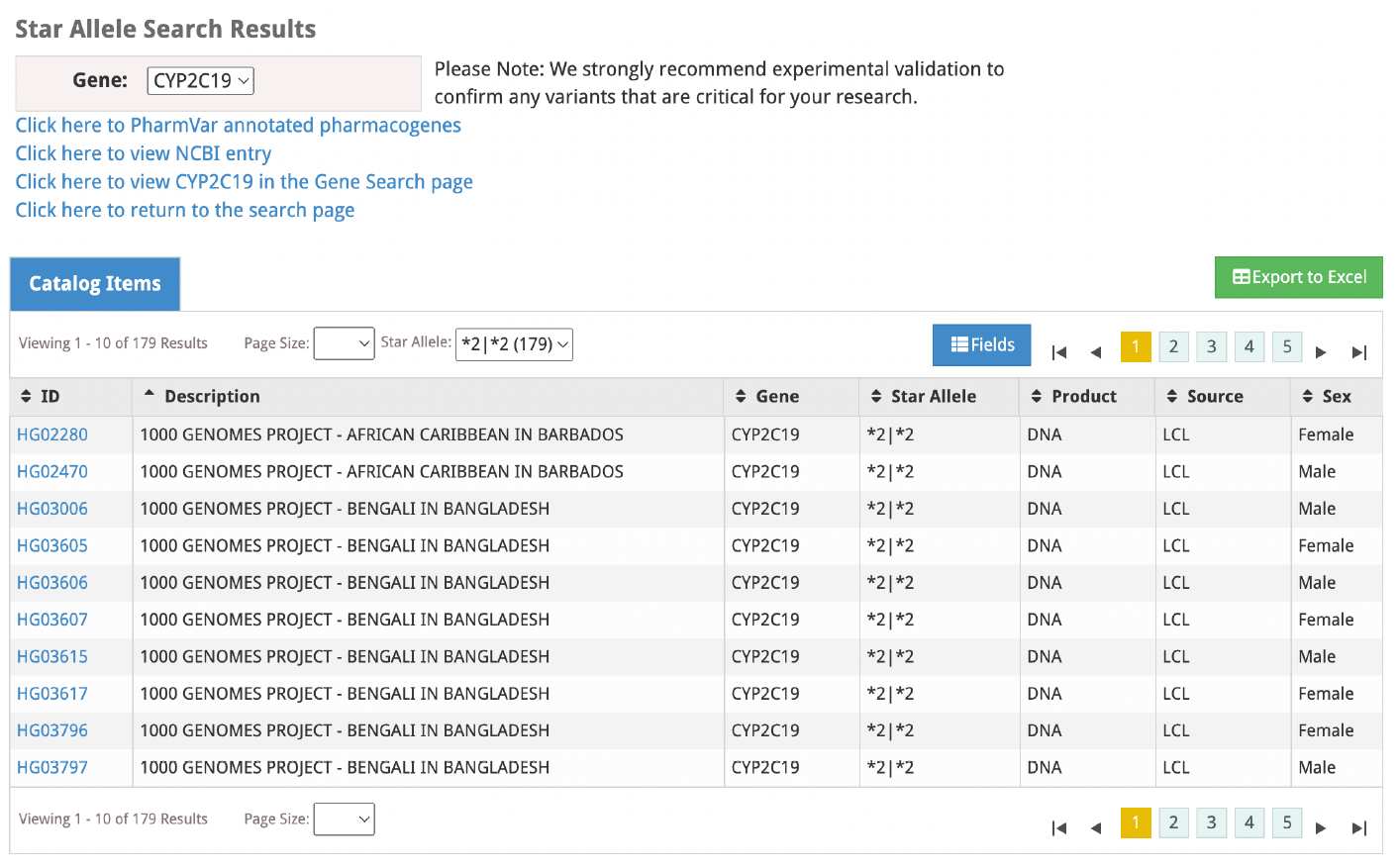
Star Allele database search results example for *CYP2C19*. Figure 2 displays a screen shot of the web-based Star Allele Search. This example is displaying results for *CYP2C19*, chosen from the dropdown search on the top, left-hand side of the page. The user may choose to view the list of PharmVar annotated pharmacogenes, the NCBI entry for the selected gene, the associated Gene Search page (which will display all of the variants included in the 30x WGS dataset for the selected gene), or to return to the general search page. The user may choose to export the Star Allele search results to a CSV file by clicking the green button on the right-hand side of the page. The user may additionally choose to filter by a given Star Allele diplotype, and this filtered drop down also displays the number of samples with each corresponding diplotypes. Figure 2 displays results after filtering for *2|*2 diplotypes in the database.

All of these genomic data search tools are designed to complement each other to ensure researchers have a simple way to search a large collection of biospecimen genetic, genomic, and transcriptomic profiles with a web-based interface that does not require bioinformatic skill or experience. For example, a researcher interested in developing a *CYP2C19* assay could first view, sort, filter and/or download a .CSV file of all of the *CYP2C19* variants included in the WGS dataset with a single HUGO symbol search (https://www.coriell.org/SNPSearch/WGS) to confirm the variants of interest are present in the data; then view, sort, filter and/or download a .CSV of the annotated *CYP2C19* *alleles for the entire sample set with Star Allele Search (www.coriell.org/StarAllele/Search) to identify the biospecimens with the relevant diplotypes; and if an alternate annotation scheme is needed (i.e., not PharmVar), the researcher can view, sort, filter and/or download a .CSV of up to 100 individual *CYP2C19* variants at a time to investigate any alternate combination of variants needed for the alternative annotation scheme (https://www.coriell.org/SNPSearch/WGS).

It is important to note all of the limitations of our approach. Our database annotations are based on short read, 30x coverage WGS and computational phasing [27]. Any error in variant calling or missing variation as well as any error in phasing in the input VCF will propagate into annotation errors. In addition, any error or missing variation in the input BAM files used for *CYP2D6* annotation will similarly produce annotation errors. While this is the most robust dataset available for our sample set at present, we anticipate that as long-read sequencing becomes more affordable and more accessible, that phase uncertainty (particularly for rare variants) will significantly go down and structural variation resolution will significantly improve. We also employed PharmVar annotation for our database and chose a strict matching requirement for each *allele annotation. This choice resulted in several ambiguous biospecimen calls in cases where one or both phased haplotypes were not an exact match to any PharmVar defined *allele.

## Conclusion

We have developed a public resource of PGx annotation for a large (n=3202) and diverse set of 1000 Genomes Project LCLs and DNAs that are available for general research use. This new resource includes a database of star allele annotation for each biospecimen and an accompanying web-based search tool (www.coriell.org/StarAllele/Search). This new tool is especially relevant to researchers interested in in vitro functional characterization of *alleles as well as for use in support of clinical PGx assay development, validation, and implementation.

## Declarations

### Ethics approval and consent to participate

All human data used in this study is publicly available through the 1000 Genomes Project [27], and all associated biospecimens belong to the NIGMS Human Genetic Cell Repository or the NHGRI Sample Repository for Human Genetic Research. All NHGRI Sample Repository for Human Genetic Research biospecimens have been consented for general research use and associated public genomic data sharing. All NIGMS Human Genetic Cell Repository biospecimens included in the 1000 Genomes Project and thereby the current study have been consented for general research use and associated public genomic data sharing.

### Consent for publication

Not applicable

### Availability of data and materials

The detailed database content is available in **Table S1** and is available through a web-based search tool (www.coriell.org/StarAllele/Search). The majority of associated 1000 Genomes Project biospecimens (LCLs and DNA) are available through the NHGRI Sample Repository for Human Genetic Research (https://catalog.coriell.org/1/NHGRI), and the collection of Utah Residents (CEPH) with Northern and Western European Ancestry biospecimens are available through the NIGMS Human Genetic Cell Repository (https://catalog.coriell.org/1/NIGMS).

### Author’s Contributions

LS designed and implemented the project and contributed to writing and editing the manuscript; NG designed the project and contributed to writing and editing the manuscript. DK, JM, and GC contributed to the design of the project and contributed to writing and editing the manuscript.

## Acknowledgments

We thank Coriell’s IT team, and John Witherspoon in particular, for supporting the implementation of Star Allele Search.

